# Deep Learning Glioma Grading with the Tumor Microenvironment Analysis Protocol for A Comprehensive Learning, Discovering, and Quantifying Microenvironmental Features

**DOI:** 10.1101/2023.06.13.544739

**Authors:** M. Pytlarz, K. Wojnicki, P. Pilanc, B. Kaminska, A. Crimi

## Abstract

Gliomas are primary brain tumors that arise from neural stem cells or glial precursors. Diagnosis of glioma is based on histological evaluation of pathological cell features and molecular markers. Gliomas are infiltrated by myeloid cells that accumulate preferentially in malignant tumors and their abundance inversely correlates with survival, which is of interest for cancer immunotherapies. To avoid time-consuming and laborious manual examination of the images, a deep learning approach for automatic multiclass classification of tumor grades was proposed. Importantly, as an alternative way of investigating characteristics of brain tumor grades, we implemented a protocol for learning, discovering, and quantifying tumor microenvironment elements on our glioma dataset. Using only single-stained biopsies we derived characteristic differentiating tumor microenvironment phenotypic neighborhoods. A challenge of the study was given by a small sample size of human leukocyte antigen stained on glioma tissue microarrays dataset - 203 images from 5 classes - and imbalanced data distribution. This has been addressed by image augmentation of the underrepresented classes. For this glioma multiclass classification task, a residual neural network architecture has been adapted. On the validation set the average accuracy was 0.72 when the model was trained from scratch, and 0.85 with the pre-trained model. Moreover, the tumor microenvironment analysis suggested a relevant role of the myeloid cells and their accumulation to characterize glioma grades. This promising approach can be used as an additional diagnostic tool to improve assessment during intra-operative examination or sub-typing tissues for treatment selection, despite the challenges caused by the difficult dataset. We present here the distributions and visualizations of extracted tumor inter-dependencies.

**Graphical abstract:** 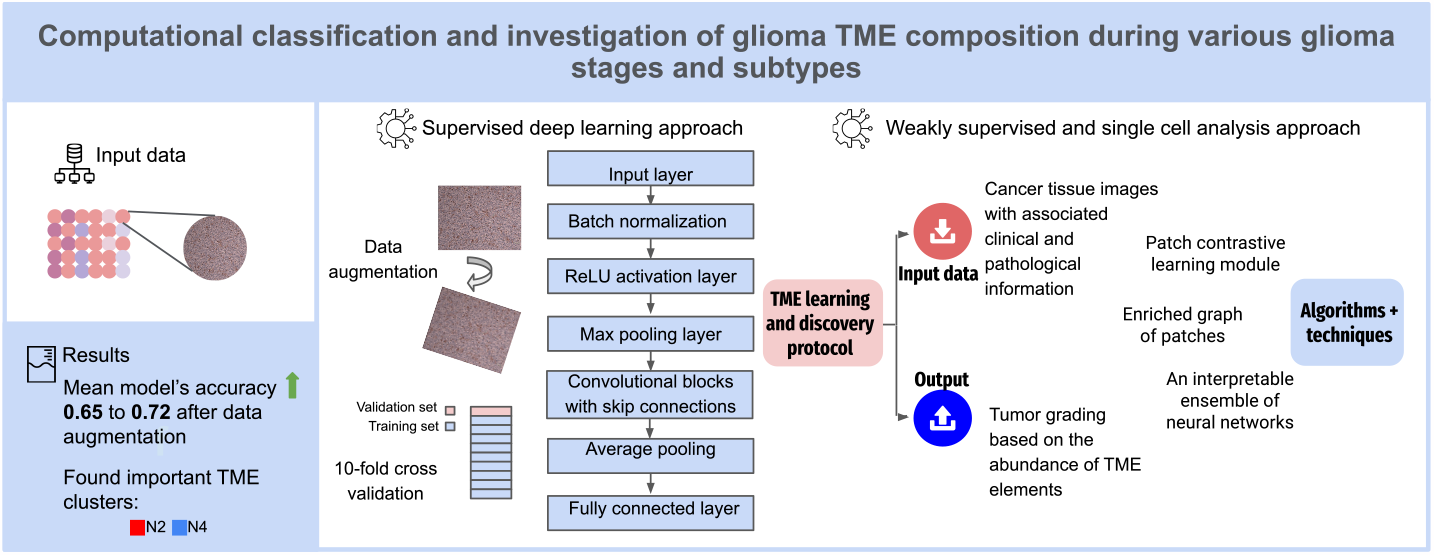

**Highlights:**

- Research highlight 1: We demonstrate that the ResNet-18 architecture with simple data augmentation trained in 10-fold cross-validation performs the multiclass classification relatively well even with a small imbalanced dataset with a high degree of similarities between classes.
- Research highlight 2: After supervised subtyping of the tumor, we investigated the usefulness of the protocol for discovery and learning tumor microenvironment elements for the same task. The protocol designed for deriving new biomarkers based on multiplex stained histological samples proved the ability to detect features characteristic of malignant tumors based only on single target stained tissue microarrays. We propose further studies on this topic can help in formulating specific criteria for improvements in diagnosis of gliomas, allowing to avoid the necessity of conducting advanced histopathological analysis or complementing genetic testing of tumor samples.

## 1. Introduction

Glioma is a primary brain tumor and glioblastoma (GBM) is among the deadliest types of tumors in adults with low survival rates even after surgery, radiotherapy, and chemotherapy in both men and women [5]. Glioblastomas are heterpgenous and comprise tumoral, stromal and immune cells that dynamically respond to many environmental stressors and therapies. In GBMs up to 30-40% of tumor mass is composed of resident and infiltrating myeloid cells [2] that participate in creating a specific tumor microenvironment (TME) supporting tumor progression. The current World Health Organization (WHO) classification criteria for gliomas combine histological and genomic indicators [36]. WHO defined three main groups of adult-type diffuse gliomas – Astrocytoma, IDH-mutant, Oligodendroglioma, IDH-mutant, and 1p/19q-codeleted, Glioblastoma, IDH-wildtype - classified according to 4 histopathological grades [23] considering tumor malignancy, nuclear atypia, and degree of invasion of the surrounding brain parenchyma. The tumor microenvironment is crucial to the tumor progression and can be manipulated to delay its progression with immunotherapy exploring various immunotools or viruses to kill tumor cells or reinvigorate anti-tumor immunity. Clinical trials are currently evaluating the safety and efficacy of an arsenal of immunotherapeutics for the treatment of newly diagnosed and recurrent high grade gliomas. The challenge is to identify which patient populations will benefit most from these medications and why [30]. Differences in immune cell proportions and phenotypes within tumors appear to be determined by glioma cell molecular characteristics [47, 52, 32]. Growing evidence highlights the significance of interactions between myeloid cells and glioma cells that enable them to evolve in a way that supports tumor growth. Spatial interactions between tumor and immune cells have been linked to clinical responses to immunotherapies in GBM patients [27].

Even if invasive, histopathology continues to be the primary method for determining the diagnosis and prognosis of cancer. In practice, a neuropathologist would investigate stained tissue slices under a microscope to diagnose and classify brain tumors.

The use of virtual slides in histopathologic analysis has expanded thanks to developments in digital microscopes collecting take high-resolution pictures of full-slide tissue specimens and tissue microarrays. Large-scale digital slide libraries like The Cancer Genome Atlas (TCGA) have made it possible for academics to access public databases of pathology images associated with clinical outcomes and genetic information. This has led to extensive research on artificial intelligence for digital pathology [10] exploring image features and quantitative classification obtained from that data to integrate different types of clinical and molecular data aiding prognosis. Mobadersany et al. [31] accurately forecasted cancer outcomes based on histology and genomics data utilizing deep learning techniques comprising convolutional layers trained to predict image patterns associated with survival. In another study, a deep learning architecture was used to create maps of tumor-infiltrating lymphocytes [40] through segmenting hematoxylin and eosin (*H&E*) stained images from the TCGA archive.

Tumor subtyping and grading is of great interest in different types of cancer [16]. In neurooncology magnetic resonance imaging (MRI) dominates in analyses of gliomas and brain metastasis [38]. In this study, we describe a similar approach but focused on histological images with human leukocyte antigen (HLA) staining highlighting immune components. We consider supervised deep learning, weakly supervised deep learning (WSDL) and single cell analysis (SCA). WSDL is built by classifying a whole slide which blindly extract tumor features in the whole slide, the latter is more focused on microenvironmental features present in patches of the image in examination and require more advanced and time consuming analysis. Those two approaches differ also in the way they are supervised as the first is given labels during the training, while the second relies on the structure and patterns within the data itself to learn meaningful representations.

The purpose of this study follows the hypothesis that automatic grading and subtyping from single stained histological slides can be an interoperative step during surgery as well as other non-invasive therapy planning [3]. Moreover, researchers anticipate that neuropathologists and machine learning models would make different sorts of categorization errors, and that the aggregate assessment of a hybrid pathologist/machine learning model will be superior to either human or machine assessment alone [26].

### 1.1. Tissue microarrays

Data used for the model training were digital images of tissue samples stained to reveal human leukocyte antigen HLA-DP, -DQ, -DR on human glioma tissue microarray (TMA) [24]. HLA antibody recognizes a major histocompatibility complex (MHC) class II heterodimer cell surface receptor expressed primarily on expressed primarily on antigen presenting cells such as B lymphocytes, monocytes, macrophages. The MHC class II molecules bind intracellularly processed peptides and present them to T-helper cells. Sample preparation is described in the next session. HLA staining reveals predominantly glioma-infiltrating myeloid cells that accumulate in the tumor and have been demonstrated to play an important role in glioblastoma biology and can be used as a diagnostic biomarker [12]. Fan et. al showed increased HLA-DR score in more aggressive stages of glioma. Authors state that HLA-DR can be used to predict the tumor grade in gliomas [9]. The choice of TMAs is justified by the fact that commercial TMAs constitute multiple, reproducible and histologically well-verified sections of tumor and normal brain tissues. Tissue microarray technique enables highly detailed immunohistochemical characterization of huge numbers of cancer cases. The use of antibodies can clearly illustrate the heterogeneity of malignant tumors [34]. Tissue sections obtained from TMA can be subjected to a vast array of regular stains, special stains, immunostainings, and morphology-based molecular methods with remarkably high-quality results [48]. TMAs are more reproducible than tumor samples or independently collected tumor sections, especially when obtained from separate laboratories. Less reproducible sections would have an impact on immunocytochemical staining and make multi-section comparison more difficult. Because TMAs were employed by so many researchers, the histopathological examination is performed by numerous pathologists, resulting in a more trustworthy evaluation. Obtaining a number of normal brain sections for comparison is usually challenging, but in the case of TMA, such sections are provided.

Novel machine learning models have emerged that outperform the typical deep learning architectures used in cancer diagnoses, such as convolutional neural networks (CNNs). While other architectures applied in the field of computational pathology include recurrent neural networks, or autoencoders [46]. CNNs have been successfully used to classify histology slides, mostly breast histology slides [17, 50, 33, 1, 29] possibly due to the availability of breast specimen histology datasets and the relative frequency of breast tumors compared to those of other organs. Although, there are studies in which authors use transfer learning to apply weights learned on one organ’s histology dataset to another organ’s dataset [21]. Ertosun and Rubin used a combination of two CNNs to categorize histology slides of glioma brain tumors into distinct tumor grades [8]. While ResNet and Inception V3 represent modern CNNs, vision transformers (ViTs) are beginning to challenge them in computer vision for natural images, indeed, Inception V3 was identified as the best-performing model [45] in image recognition methods across five public histopathology datasets. On the other hand, Chen et al. [6] implemented a ViT leveraging the natural hierarchical structure inherent in whole slide images and it outperformed other models for cancer subtyping. However, transformer models lack the translation equivariance and locality of CNNs; as a result, vision transformer is less effective with limited datasets. They are computationally heavy (especially compared to CNNs) due to their self-attention mechanism, memory-intensive having a large number of parameters, and have limited generalization on small datasets.

Lastly, by using transfer learning to pre-train a ViT on large-scale datasets significantly enhances its accuracy [25]. Apart from supervised deep learning that requires annotated data, there are two primary computational pathology strategies for automating the analysis of the histopathology or phenotype of a tumor: weakly supervised deep learning (WSDL) which automatically infers fine-grained (pixel- or patch-level) information from coarse-grained (image-level) annotations, and single cell analysis (SCA) which by segmenting the cells in the tissue and measuring their shape and marker expression intensity identifies clusters of cells with similar characteristics, as well as higher-order interactions or phenotypic ’neighborhoods’ [42] are additional methods for automated histology analysis. The objective of weakly supervised learning is to automatically infer fine-grained (pixel-or patch-level) information from coarse-grained (image-level) annotations. With all these advancements, a pathologist’s annotation workload is drastically reduced by weakly supervised learning [49, 46].

SCA methods [41] start by segmenting the cells in the tissue and measuring their shape and marker expression intensity. This data is then utilized to identify clusters of cells with comparable characteristics, as well as higher-order interactions or phenotypic ’neighborhoods’ [42]. SCA techniques accomplish this by constructing topological networks with cell phenotypic interactions and using graph-based clustering to allocate groups of cells to distinct neighborhoods. SCA approaches give a high level of interpretability since they use the cell as the main unit of tissue depiction [4]. The cell-to-graph approach extracts individual cells from the images and represents their spatial relationships as graphs allowing tracking the intricate cellular interactions within tumors. In tumor grading, the cell-to-graph approach can be used to automatically quantify various morphological features of cells, such as size, shape, and arrangement, and classify tumors into distinct grades. By representing cells as nodes and their spatial relationships as edges in a graph, the cell-to-graph approach enables the analysis of cellular interactions, such as cell clustering, spatial distribution, and connectivity patterns, which can provide valuable insights into the behavior and prognosis of tumors. This approach can help automate and standardize tumor grading, reduce subjectivity and inter-observer variability, and provide quantitative and reproducible measures for tumor characterization, which can aid in clinical decision-making.

Zhou and colleagues [53] created a cell graph convolutional network utilizing nuclear appearance features and spatial location of nodes to enhance its performance. Cell graph convolutional network also introduces Adaptive GraphSage, a graph convolution technique that combines multi-level features in a data-driven manner, to enable nodes to fuse multi-scale information. To address redundancy in the graph, a sampling technique is proposed to remove nodes in areas of dense nuclear activity. The cell graph convolutional network approach can capture the complicated tissue microenvironment by considering images 16 times larger than patch-based methods. Pati [35] introduced a novel multi-level hierarchical entity-graph representation of tissue specimens for cancer diagnosis and prognosis. Traditional approaches have encoded cell morphology and organization using cell-graphs, but this study proposes a hierarchical graph representation that captures histological entities at multiple levels, from fine to coarse. Study demonstrates the potential of using multi-level hierarchical entity-graph representations for encoding tissue structure and function in cancer diagnosis and prognosis, and it is evaluated on a large cohort of breast tumor specimens. Lastly, there have been works on explainable graphs in digital pathology. This refers to graph-based techniques that are employed to represent and assess digital pathology images in a way that provides more control over concept representation and allows for comprehensive and compact explanations. Jaume et al. [15] introduced a post-hoc explainer that derives compact per-instance explanations emphasizing diagnostically important entities in the graph. They prune the redundant and uninformative graph components, and the resulting sub-graph is defined as the explanation. Yu et al. [51] proposed a method called IFEXPLAINER, which generates necessary and sufficient explanations for graph neural networks. The explanations are based on the measurement of information flow from different input subgraphs to the predictive output of the network. The authors introduced a novel notion of predictiveness called f-information, which incorporates the realistic capacity of the graph neural network model, and use it to generate the explanatory subgraph with maximal information flow to the prediction. The produced explanation is essential to the prediction and sufficient to reveal the most crucial substructures.

Taking into consideration the lessons learned and limitations of the aforementioned approaches, we investigated the two scenarios of whole slide classification and neighborhood of single cells analysis, which are described in the following sections.

## 2. Methods

### 2.1. Data

The tissue microarrays used in the experiments are complex, commercially available paraffin blocks constructed by extracting cylindrical core tissue ”biopsies” from different donor paraffin blocks and embedding them again in a single recipient block (microarray) at specific matrix coordinates [48]. Of those TMA were sections representing 21 normal brain tissue samples, 24 samples of WHO grade 1, 84 – grade 2, 26 – grade 3, and 48 – grade 4 gliomas (Fig. 1). 63% of malignant samples were collected from male patients and 37 % of malignant samples – from female patients. For the sake of convenience and consistency, we named normal tissue samples “grade 0”.

**Figure 1.**
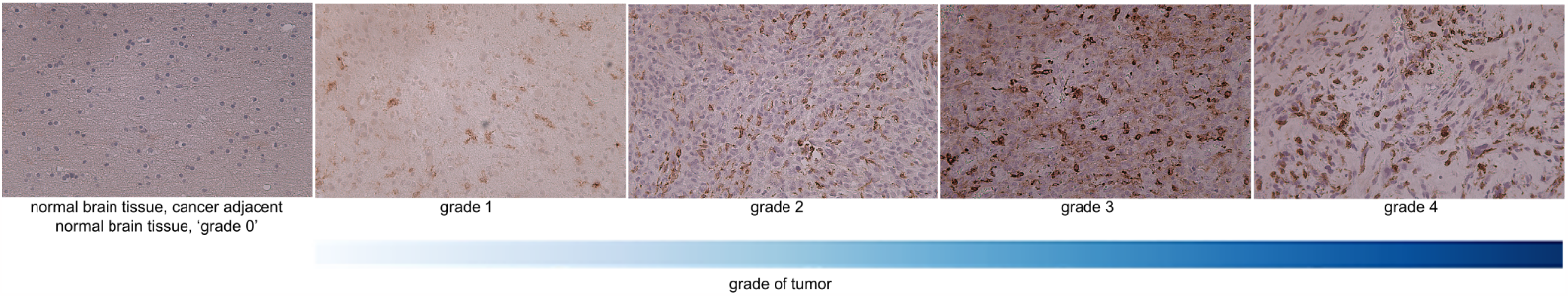
Examples of normal brain tissue and *H&E* stained tissue microarray core images of each glioma grade.

### 2.2. HLA immunostaining

We have used commercial GL2083b and GL806g TMAs from TissueArray.LLC.com (MD, USA) containing multiple GBM cases, normal brain tissues with duplicate cores per case. Paraffin-embedded sections were incubated 10 minutes in 60°C, deparaffinized in xylene, rehydrated in ethanol (100%, 90%, 70%) and washed with water. Epitopes were retrieved by boiling in a pH 6.0 citrate buffer for 30 min. Further steps (blocking of endogenous peroxidase and un-specific interactions, application of secondary antibodies and signal developing) were performed using reagents from ImmPRESS® kit according to the manufacturer’s protocol. Primary antibody recognizing HLA-DP, -DQ, -DR antigen (anti-HLA-DP, -DQ, -DR, Dako, #M0775) diluted in 3% donkey serum was applied for overnight incubation in 4^°^C. All washes were performed using PBS-T (Tween-20, 0.05%). Sections were counterstained with hematoxylin washed in H2O, dehydrated and mounted using a glycerol mounting medium. Images were acquired with the Leica DM4000 B microscope operating with the Application Suite ver. 2.8.1 software (Leica Microsystems CMS, Switzerland).

### 2.3. Supervised deep learning approach

Small sample size and unbalanced data distribution posed two main challenges for the subsequent multiclass classification task; as images of WHO grade 2 tumors were approximately three times overrepresented in comparison to other grades. To address the issue of imbalanced dataset and check if the model’s performance will improve we split in a cross-validation style for the test and training subsets, and then augment histologies of normal brain tissue and WHO grades 1, 3, 4 by rotating the view of 90 and 180 degrees. As most of the deep learning architecture are not rotation invariant rotated images can be considered as separated samples [14].

During preprocessing the tissue microarray well views were center cropped. In the phase between the preprocessing and further experiments, we investigated the impact of removing difficult image cases that showed significant discrepancies within classes in our TMA dataset. We found that the overall accuracy did not improve substantially after removing 9 such instances. Therefore, we decided to omit the step of cleaning an already limited dataset from images of cores with a lower amount of tissue. We will present results for the case of training with unbalanced small sample size, as well as an augmented dataset with a more balanced distribution.

We chose to use a CNN architecture not susceptible to vanishing or exploding gradients, which is addressed by a deep residual learning framework [11]. In this study, the model implemented was the simplest among ResNets - ResNet-18, with a 7×7 convolutional layer filter, then 16 convolutional layers with 3×3 size filter and average pooling. Output was passed through SoftMax activation function. The model was trained within 10 iterations for different data shuffle to split subsets and augmentation. Data augmentation was performed on the training dataset avoiding redundancy in the validation set. We conducted 2 experiments: 1) training ResNet-18 from scratch and 2) applying a pre-trained model and updating only the last layer’s weights.

### 2.4. Single cell analysis & weakly supervised deep learning

In the second part of our study, we expanded a previously introduced pipeline for cancer tissue analysis [18], with the aim of quantifying the glioma microenvironment. The approach integrates cell-level interpretable quantification of the TME with patch-based weakly supervised learning of tumor histopathology to automate the in situ discovery of clinically relevant TME elements. The learning and quantifying protocol is a multi-level, interpretable deep learning ensemble that discovers the most important TMEs from tissue sections while performing a classification task using only patient-level labels. The algorithm assigns patches to TMEs at three levels of spatial complexity: local cell phenotypes, cellular neighborhoods, and tissue area-specific interactions between neighborhoods. The protocol was initially designed for retrieving computationally specific cancer biomarkers from multiplexed stained tissues - here we applied The learning and quantifying algorithm to search if a useful portion of interactions and abundances of tumor features along grades of malignancy could be retrieved. The input data consists of images of HLA-DR-stained (single stain) glioma TMAs with clinical and pathological data on their WHO grades. The patch contrastive learning module divides images into patches and embeds each patch in a 256-dimensional vector using a CNN that has been trained unsupervised to assign similar vectors to patches containing similar biological structures.

Then, a graph of patches containing the spatial interactions between tissue patches is generated. The learning and quantifying algorithm is an interpretable ensemble of neural networks that learns phenotypes, phenotype neighborhoods, and areas of interaction between neighborhoods in order to classify patients based on the abundance of TMEs. Implementation discovers tumor landscape characteristics that are relevant to a certain predictive task. This can be done a posteriori by analyzing the TMEs obtained by the ensemble of networks while classifying patients. It is the purpose of the BioInsights module, which is accomplished by identifying cohort-differentiating features (differential TME analysis) and significant TMEs in individual predictions (predictive influence ratio - PIR). Codes of the methods described above are provided: https://github.com/octpsmon/TME_analysis_protocol_n_glioma_grading.

## 3. Results

### 3.1. Results from supervised deep learning classification

The best trained-from-scratch model’s accuracy achieved a value of 0.77 and the mean accuracy from 10 iterations is 0.72 with 0.03 of standard deviation on the validation set. Training with data augmentation provided better results than the one without regularization (0.65 of mean accuracy with 0.06 of standard deviation (SD) and the best score achieved at level 0.76). Model average scores compared in Table 1 demonstrate that simple data augmentation improved the efficacy of the network. Pre-trained ResNet-18 achieved better mean accuracy than the model trained from scratch as depicted in Table 1. Without data augmentation mean validation accuracy was 0.83% with SD of 0.08. When using data augmentation regularization technique the mean validation accuracy was 0.85% with a SD of 0.05. The best score from k-fold cross-validation with and without data augmentation applied was equally 0.92.

**Table 1:**
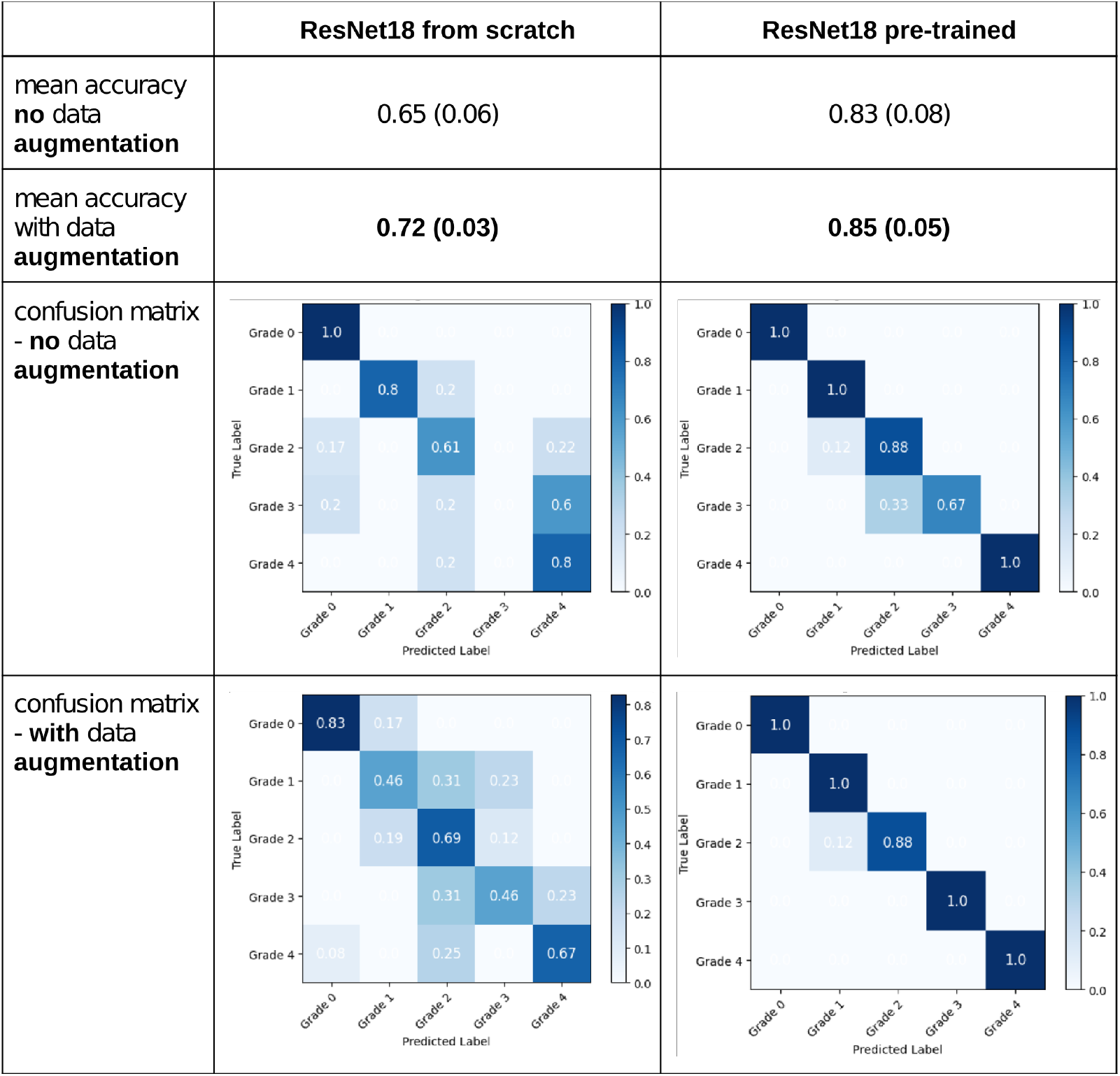
Table presenting results of training ResNet-18 model from scratch with (left) and updating only last layer’s weights using model pre-trained on ImageNet dataset (right). The table compares models’ validation accuracies when they were trained with and without data augmentation. The table includes normalized confusion matrices that visualize the performance of an algorithm, representing the instances in an actual class and the instances in a predicted class.

|  | ResNet18 from scratch | ResNet18 pre-trained |
| --- | --- | --- |
| mean accuracy<br>no data<br>augmentation | 0.65 (0.06) | 0.83 (0.08) |
| mean accuracy<br>with data<br>augmentation | **0.72 (0.03)** | **0.85 (0.05)** |
| confusion matrix<br>- no data<br>augmentation |  |  |
| confusion matrix<br>- with data<br>augmentation |  |  |

### 3.2. Results from weakly supervised deep learning and single cell analysis

Results from contrastive learning reveal the model has strong discriminatory power (false positive rate of 0,8 - 0,9) to differentiate between grades 0 and WHO grade 1, as well as 1 and WHO grade 4. The model obtained good results (false positive rate of 0,7 - 0,8) when distinguishing between the following grades: (0, 4), (1, 2), (1, 3). An acceptable score (false positive rate of 0,6 - 0,7) was shown in the discrimination between grades: (0, 2), (0, 3). Patch contrastive learning is not enough reliable when it comes to differentiating grades: (2, 3), (2, 4), (3, 4) (Fig. 2).

**Figure 2.**
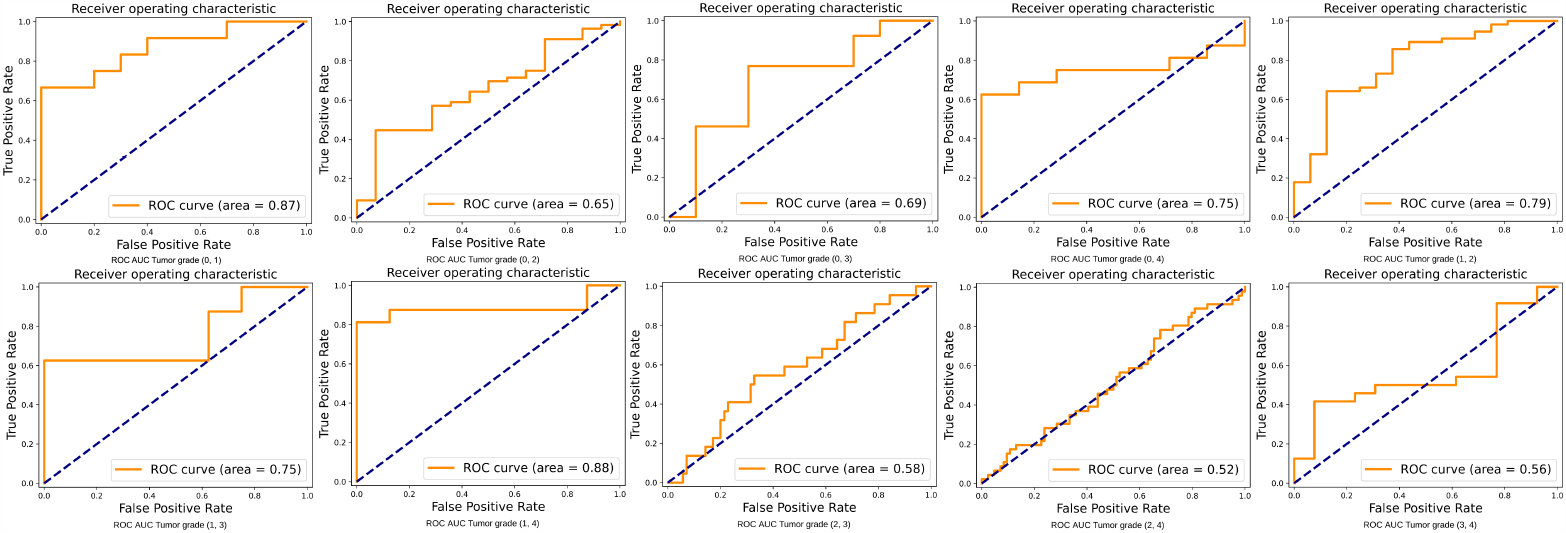
Comparison of receiver operating characteristic (ROC) area under the curve (AUC) for tumor grade classification across 10 comparisons.

The learning and quantifying algorithm learned to recognize 23 patterns of TME elements - 8 phenotypes, 8 cell neighborhoods, and 7 areas of neighborhoods. The quantitative model distinguishes the highest number of differences between grade 0 and WHO grade 4. It also recognizes a high significantly increased abundance of cells of neighborhoods N2 in WHO grade 4 compared to WHO grade 1 as well as a minimally significantly increased abundance of the neighborhoods N4, N6, N8, and N9 of cells and phenotypes P3, and P4 comparing these two. Neighborhood N2 detected in WHO grade 2 glioma is less significantly increased compared to grade 0 than in WHO grade 4 compared to 0. Therefore, N2 is highly predictive for patients with tumor of WHO grade 4. The neighborhood N4 appears significantly more frequently in WHO grade 3 compared to grade 0, suggesting that N4 was relevant to successfully classifying WHO grade 3. The neighborhood N3 is visible significantly more often in WHO grade 4 compared to WHO grade 2.

N2 occurs most frequently on TMA core images (Fig. 4) and its abundancy has the highest significance when comparing tumor grade 0 with WHO grade 4, significance of p-value less than 0.01 when comparing WHO grades 1 with 4, and significance of p-value less than 0.05 when comparing grade 0 with WHO grade 2. N9, the second neighborhood of phenotypes, is associated with tumor WHO grades 1, 2, and 4 (Fig. 6). On the relative abundance plots, we can observe in detail how the neighborhoods N2 and N4 are the two showing the most significant differences between grades (Fig. 5). Figure 3 presents the actual view of the patches assigned to clusters N2 and N4. P3 and P4 are the phenotypes that appear most frequently in images of high grade gliomas. The N4 neighborhood’s view might reflect an accumulation of protumor macrophages resulting in increased HLA staining. The extracted characteristics include round and ameboid cells.

**Figure 3.**
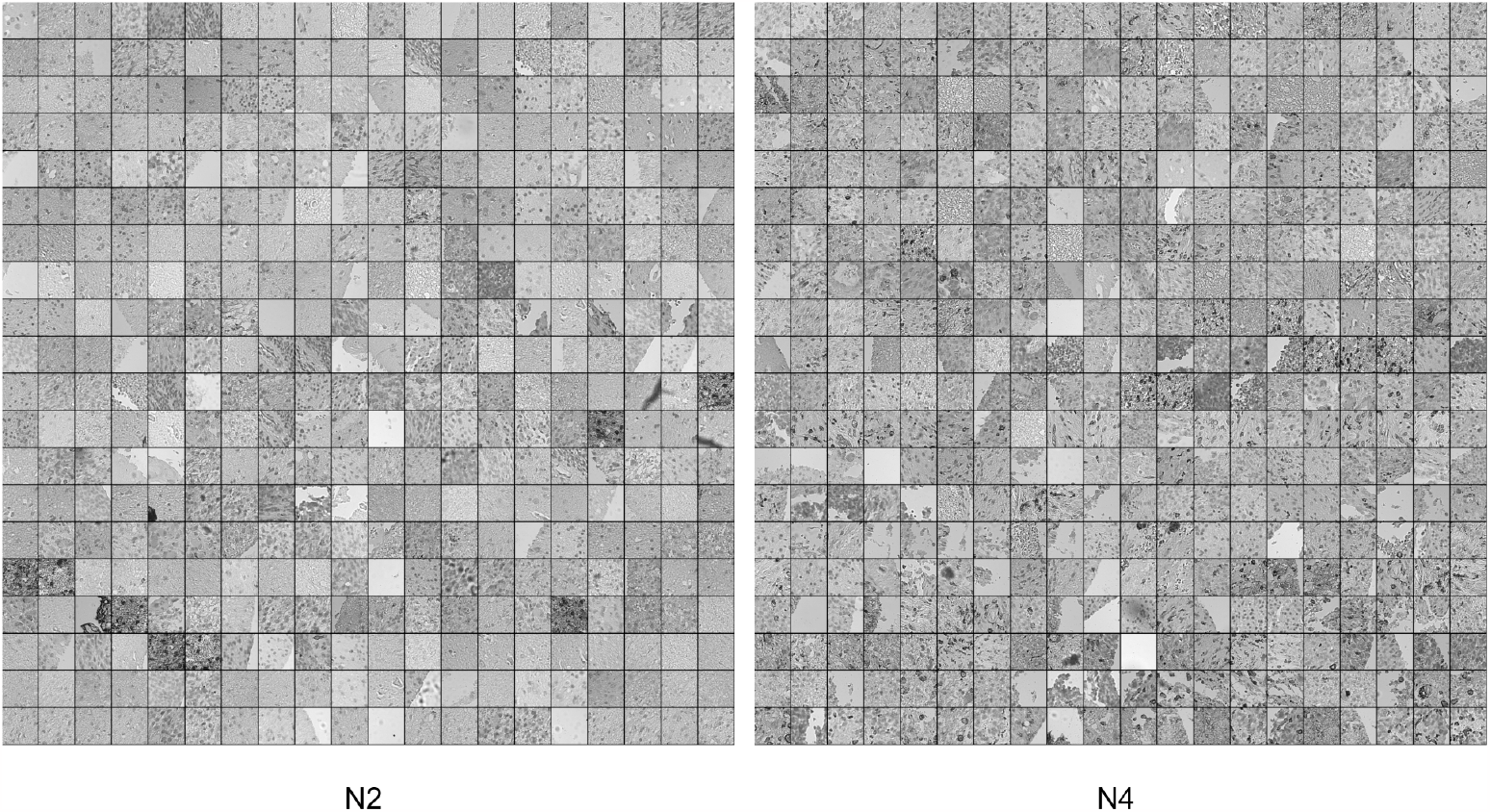
View of clusters (neighborhoods). This figure presents 2 frames made of patches of the tumor images that were assigned together within 2 groups during patch contrastive learning protocol and resulted in the 2 most distinctive clusters of cell neighborhoods.

**Figure 4.**
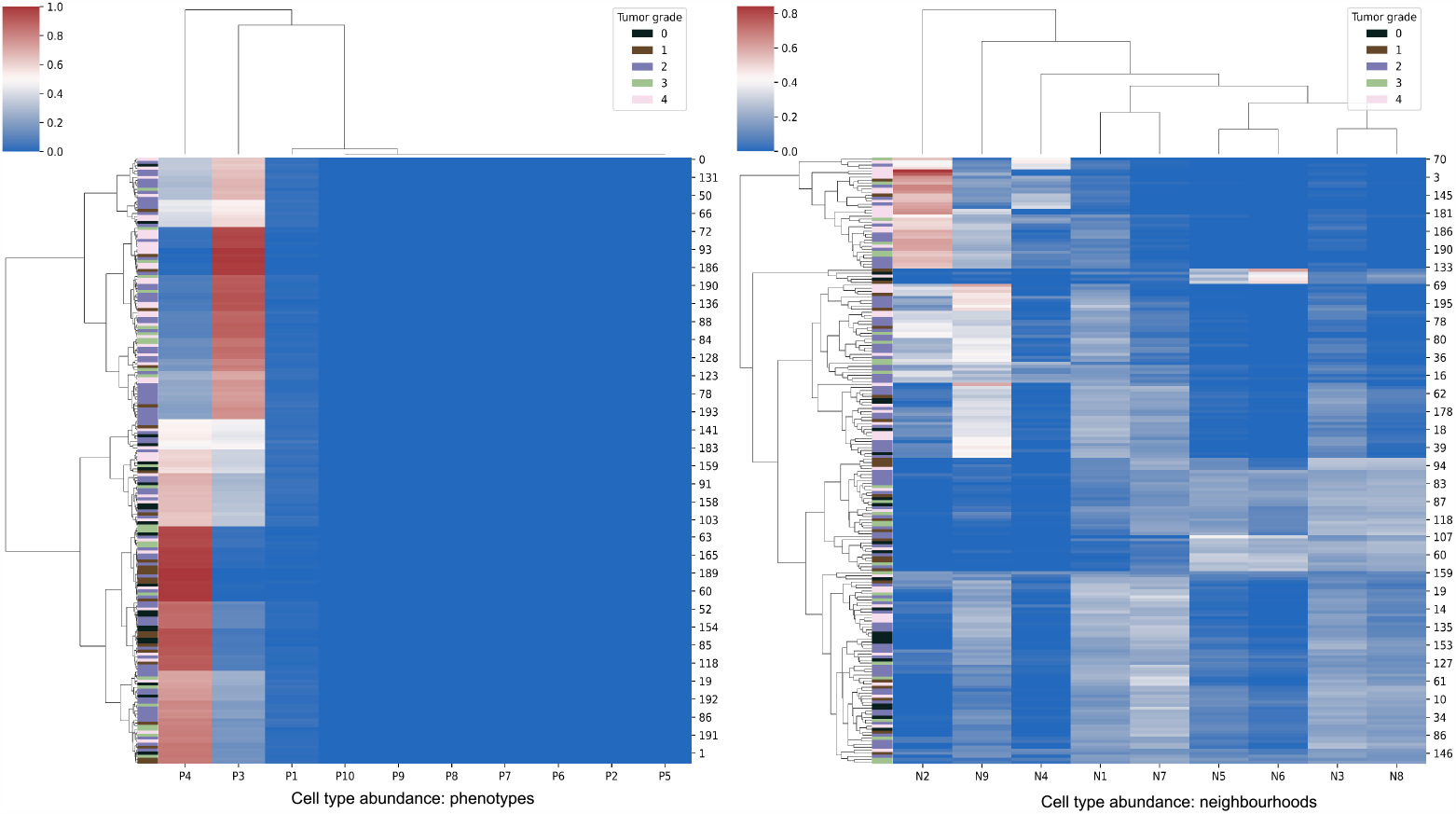
Cell type abundance. TME composition of fields - a heatmap showing the sum-to-1 normalized distribution of neighborhoods in 203 images/patients, colored by label (tumor grade). Both images/patients and neighborhoods are ordered by hierarchical clustering. The most abundant phenotypes are P4 and P3. The most abundant neighborhood is N2 - mostly in grade 4 and 2.

**Figure 5.**
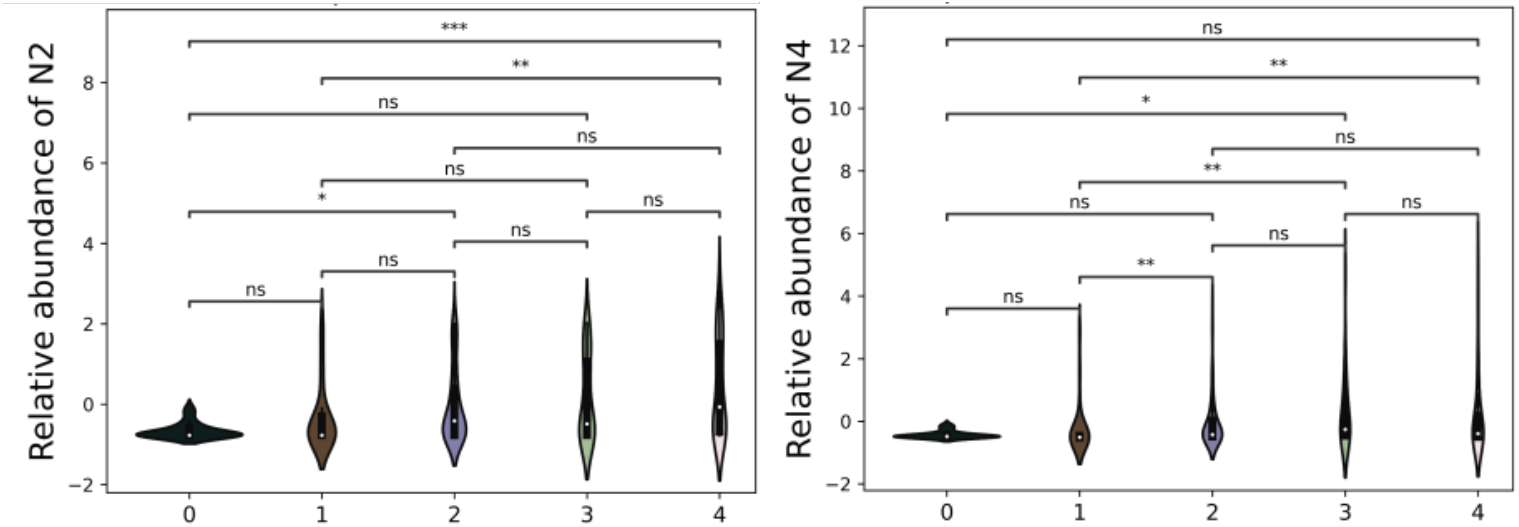
Violin plots showing a relative abundance of clusters. The plots represent how significant are particular differences between cluster abundance between tumor grades. The distribution of each plot illustrates the variation and density of the cluster abundance, emphasizing the statistical significance of group differences. Marking of *** indicates p-value of <0.001, marking of ** - the p-value of <0.01, and marking of * - the p-value of <0.01

**Figure 6.**
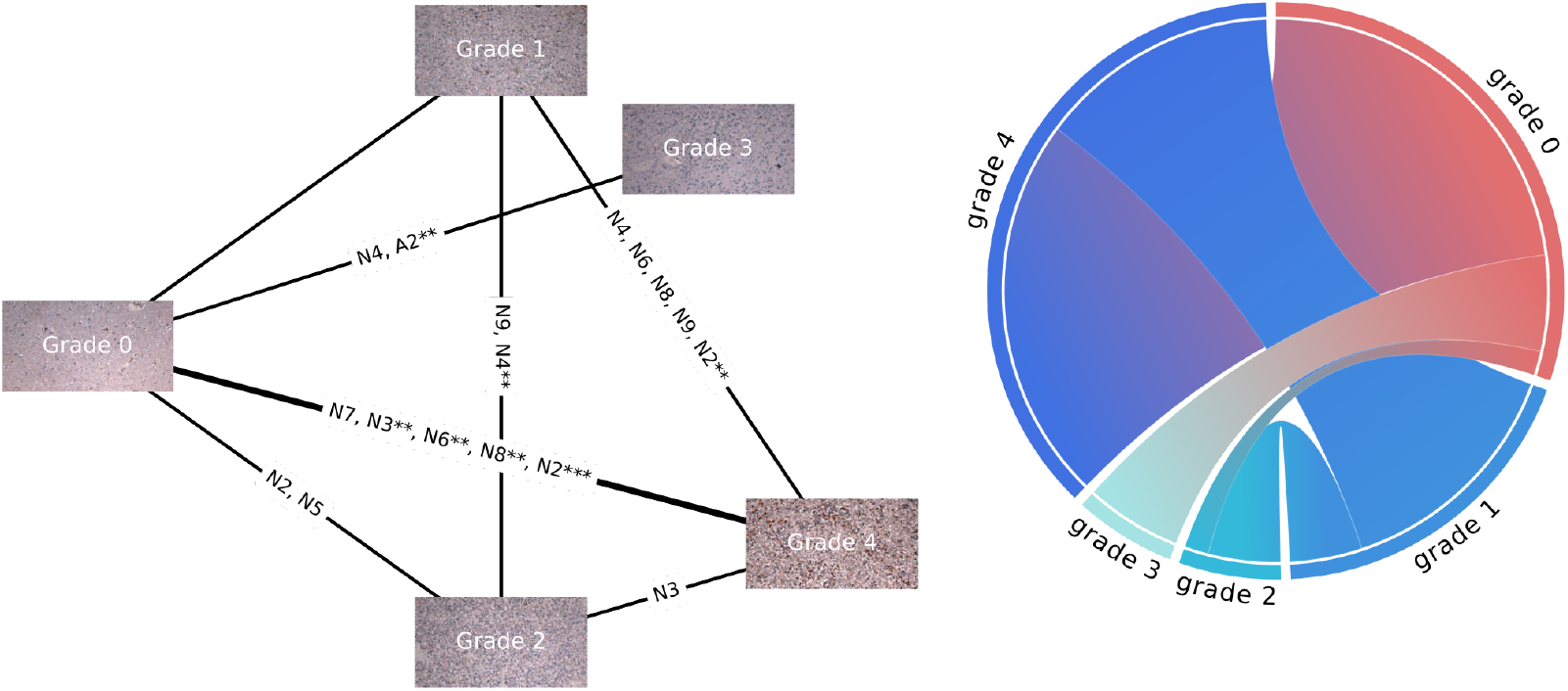
Graph of clusters of tumor microenvironment features. On the left: the undirected graph with connections linking tumors of various grades where occurred significant differences between cluster abundances. The bold line indicates two grades with the most different characteristics. On the right: a weighted sum of common clusters occurring with significant differences in the abundance between tumors of different grades. The diagram on the right is based on the same results as the graph on the left, but it more strongly represents the number and significance of the differences between grades (when the p-value is <0.01 the number of shared clusters is multiplied by 2, and when the p-value is <0.001 - the number of shared clusters is multiplied by 3).

The global differential TME analysis revealed that the highest predictive influence have el-ements: N1, N2, N5, N8, N9, which corresponds with measurements of significance of their abundance in 203 images of different tumor grades. The higher the predictive influence ratio value, the higher the relevance of the TME element for a specific patient.

Violin plots (Fig. 5) present significant differences between derived clusters’ between tumor grades. N2 neighborhood occurs highly significantly more abundantly in the most malignant tumor than in tumor adjacent tissue and moderately more rich in the WHO grade 4 rather than in 1. While N4 neighborhood appears significantly more frequently in grade 4 than in 1 tumors, but also in WHO grade 3 compared to 1. There is significance observed when comparing relative abundance of cluster N4 between grade 3 and normal brain tissue.

### 3.3. Visualization with UMAP

To further investigate our data, we applied UMAP (Uniform Manifold Approximation and Projection for Dimension Reduction) [28]. By constructing a low-dimensional embedding of high-dimensional TME and tumor grade data, UMAP revealed patterns and relationships that are challenging to discern in the original high-dimensional space (Fig. 7). Specifically, it identifies clusters of tumors with similar grades and reveals underlying substructures that may be indicative of distinct tumor subtypes. When the UMAP plotting program was applied to raw HLA images from tumors of various grades, it showed normal and tumor adjacent brain tissue as well separated from the tumor samples. WHO grade 2 tumors are the most spread among others. Tumors of WHO grade 3 and grade 4 form dense and proximate groups in the same location. UMAP plot of phenotypes shows noisy, not very informative patterns. In turn, when phenotypes are grouped into neighborhoods and only after that are applied as input for the UMAP - they reveal a well-defined cluster of the neighborhood N9 - which represents the most abundant feature on the images within glioma dataset 4, as well as concentrated N2 with the highest significance in comparisons between grades. This UMAP plot we can also consider an evaluation of contrastive learning results from the learning and quantifying algorithm protocol.

**Figure 7.**
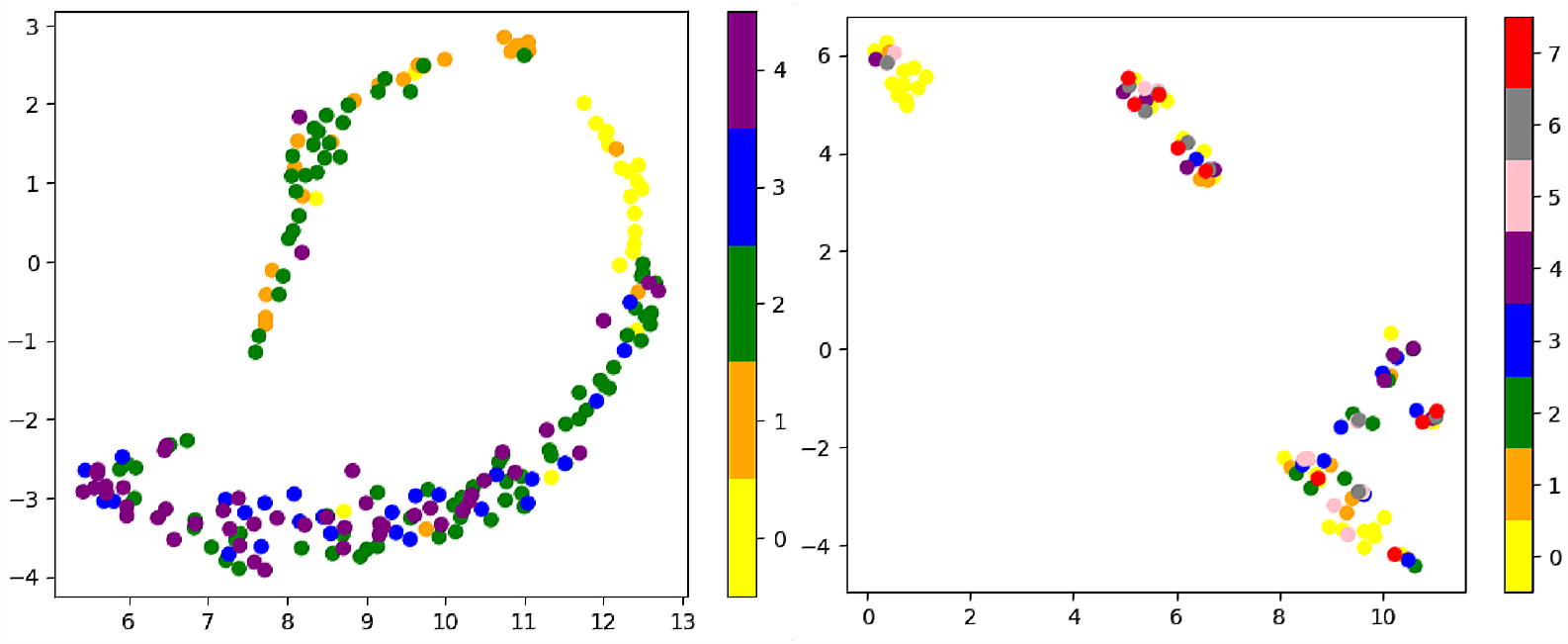
Left: UMAPs of tumor grades. Labels in the form of color-coded for tumors with specific grades are assigned as follows - starting from the left: 0 - tissue ”grade 0”, 1 - tumor grade 1, 2 - tumor grade 2, 3 - tumor grade 3, 4 - tumor grade 4. Right: UMAPs of neighborhoods: 0 - N1 neighborhood, 1 - N2 neighborhood, 2 - N3 neighborhood, 3 - N5 neighborhood, 4 - N6 neighborhood, 5 - N7 neighborhood, 6 - N8 neighborhood, 7 - N9 neighborhood.

## Discussion

In this work, we investigated automatic brain tumor grading by using supervised classification and a single cell analysis weakly supervised. The results of the supervised classification demonstrate that simple data augmentation enhanced the network’s performance. As the confusion matrix shows, the model tends to often misclassify WHO grade 2 with grade 3 tumors. This is probably related to general difficulties even for experienced histopathologists distinguishing grade 2 from 3 of those tumors, so the model behaves ‘pessimistic’, assuming a higher grade of the tumor in advance. That means it does not classify as 2nd-grade astrocytoma, oligodendroglioma, or oligoastrocytoma, but rather as an anaplastic type of those. The principal distinguishing feature of WHO grade 3 gliomas is the increased mitotic activity and histological anaplasia but the threshold mitotic count for a WHO grade 3 designation has not been established yet. WHO 2021 classification added molecular markers connected with poorer prognosis that are recommended for histological diagnosis in assigning the grades 2 or 3 [39, 22].

We performed single cell analysis with weakly supervised learning. Differential composition analysis between clusters of TME elements involves identifying differences in the composition of tumor microenvironments between distinct groups of cells or tissue clusters. The goal was to identify specific cellular or molecular features that are associated with different stages of tumor development (or could be also associated with a response to treatment). Tumor microenvironment elements include a wide range of cells, such as immune cells, stromal cells, and tumor cells, as well as extracellular matrix molecules and signaling molecules. By analyzing the composition of these elements in different groups of cells or tissue clusters, we are able to gain insights into the cellular and molecular mechanisms that drive tumor progression.

By using machine learning algorithms like the contrastive learning presented in the previous sections, we can analyze large amounts of brain imaging data more efficiently and accurately. They can identify subtle patterns of HLA-DR staining that may not be immediately apparent to the human eye, and group together brain regions that exhibit similar patterns of HLA-DR staining. This approach unveils previously unrecognized similarities and differences in the activation of glioma-associated microglia, macrophages and other immune cells in the brain, providing a more comprehensive understanding of the underlying biological processes that contribute to neurological and psychiatric conditions. The abundance of HLA-expressing myeloid cells and their acquisition of the ameboid phenotype have been demonstrated in glioblastoma [19, 44]. In turn, this information can inform the development of more targeted diagnostic and therapeutic approaches that can improve patient outcomes.

Our approach can be considered an adequate substitute for spatial transcriptomics tools when one would like to investigate cell populations in TME. There are frameworks available for tissue annotation with pixel-level resolution that integrate histological information with gene expression [13]. In this way the gene expression can be integrated with histological information directly based on the histology image. The method represents a great way to understand the spatial architecture of the TME, however, unlike in our pipeline, it requires the step of next-generation sequencing (NGS) of the tumor sample.

We demonstrate that our investigation of cell population abundances within and across distinct tumor grades revealed consistent findings that align with previous studies. False positive rates with their satisfying scores when comparing normal brain and low grade glioma to high grade glioma and poor result when comparing grade 2 and 3 correspond to the supervised model outcomes and to the clinical practice. In the evaluation of abundance significance when comparing clusters identified within TME of tumors of various grades, we did not find a significant dissimilarity between WHO grades 2 and 3. Although we measured that the mean relative abundance of N9 neighborhood in WHO grade 3 samples lays at the level 0 and at the same time - at the level of 0.5 in the case of WHO G2. This is consistent with clinical practice as a reliable dissection of WHO grades 2 and 3 tumors is possible with the inclusion of molecular markers.

Gliomas being intricate tumors with a complex interplay between tumor cells and the immune system, necessitate robust staining methods to unravel their immunological landscape. The choice of tissue staining is far from trivial in this context, as it directly impacts our ability to decipher the intricate immune response within the glioma microenvironment.

HLA-DR staining is a method used in immunohistochemistry to identify the expression of human leukocyte antigen (HLA)-DR, DP, DP, DQ in tissue samples. HLA-DR is a major histocompatibility complex (MHC) class II antigen that is expressed on the surface of antigen-presenting cells and is critical for the initiation of an immune response. HLA-DR staining can reveal the presence and distribution of myeloid cells such as microglia, monocytes, macrophages, and dendritic cells that are immunosuppressive and tumor supportive [20] The abundant expression of HLA staining and acquisition of the ameboid phenotype by HLA expressing cells have been demonstrated in high grade gliomas [19, 44]. In glioma tissue, malignant gliomas are known to modulate the immune response, leading to an immunosuppressive tumor microenvironment. MHC class II antigens, specifically HLA-DR and DQ, are critical for lymphocyte response and that transfection of these genes may boost the immunogenicity of glioma cells [43]. Fan and colleagues [9] consistently found higher mRNA expression of five HLA-DR genes shows association with a higher tumor grade and the HLA-DR score, which described the expression of all HLA-DR genes, revealed higher levels associated with a higher tumor grade. In comparison to oligodendroglial tumors, astrocytic tumors, the more aggressive histological subtype of lower grade gliomas, exhibited a considerably higher HLA-DR score. Tumors with the increased HLA-DR expression should have higher immunogenicity, lesser aggressiveness and a better prognosis. However, gliomas with high HLA-DR scores exhibited low immunogenicity, few defined tumor antigens, and more invasive features [7], and patients with high HLA-DR scores had poor survival rates in lower grade gliomas. This conflicting finding about the relationship between a high HLA-DR score and a poor clinical outcome could result from the immunosuppressive reprograming of myeloid cells in glioma TME and complexity glioma immunogenicity.

Neighbourhood N2 abundancy shows the highest significance when comparing tumor grade 0 with grade 4 which agrees with the estimation of HLA-DR scores measurements done by Fan et al. [9]. They reported lower HLA-DR mRNA levels in normal brain tissue, low grade glioma (WHO grade 1,2), and higher in anaplastic glioma (WHO grade 3), and glioblastoma (WHO grade 4). They evidenced corresponding differences including p¡0.001 between normal brain tissue and glioblastoma.

The clusters of neighborhoods N2 and N4 extracted from the model (Fig. 3) can represent pattern of the increased HLA-DR staining in WHO grade 4 gliomas indicating accumulation of activated, tumor supportive myeloid cells within the tumor core and changing the local tissue structure and vascularization. Their increased abundance and activation (reflected by more intensive HLA-staining) contribute to a greater complexity of images detected by the protocol. We present the view of the neighborhood N9 occurring more abundantly in the WHO grade 2 than in 3.

With the increasing complexity and abundance of imaging data, there is growing optimism that computational methods and advanced image analysis techniques will play a pivotal role in deriving meaningful visual features of tumor clusters. Visual inspection of extracted clusters serves insights into the spatial arrangement, cellular composition, and architectural patterns within the tumor microenvironment, providing a rich source of information for characterizing tumor heterogeneity and progression.

In the pursuit of comprehending histology classification and unraveling the intricate landscape of the tumor microenvironment, we find ourselves confronted with evolving classification criteria towards more accurate and unambiguous. The WHO adopted ”across tumor types” rather than ”within a tumor type” grading. Regardless of the histological variability, tumors of the same grade have essentially the same clinical outcome. However, the issue of inter- and intra-observer variability remains unresolved. The evaluation of atypia is subjective, while mitotic statistics are dependent on the thoroughness of the examiner as mitoses can accumulate focally or uniformly [22]. Combination of histopathological findings with molecular data allows for the more reliable classification of brain tumors, and the inclusion of the unsupervised learning analyses of DNA methylation and NGS data has identified distinct subgroups with distinct molecular characteristics [37]. However, the acquisition of DNA methylation and NGS data is expensive and not always affordable for patients, therefore the computational quantification AI-supported protocols dedicated to histology images may contribute to formulating unique descriptors of glioma subtypes complementing diagnosis.

## Conclusions

CNN based diagnostic tools are supervised or weakly supervised deep learning methods that have been effectively applied to a variety of image analyses. They require large labeled datasets, that are not always available in the medical field of rare tumors. We present a new approach dedicated to classifying gliomas, detecting malignant ones and distinguishing them from normal brain tissues based on images acquired from a single HLA-staining of tissues. Given the small sample size and high degree of similarity between WHO grades 2 and 3, this was a particularly challenging task. Even with a limited sample size, the proposed architecture for deep learning performs relatively well. The application of cell abundance analysis allowed to retrieve the distinct neighborhood of phenotypes, including the N2 that occurs most often on the TMA core images and shows a highest significance when comparing tumors of grade 0 with 4. The second characteristic (differentiating tumor grades) neighborhood of phenotypes is N4, and the second most abundant one is N9. Phenotypes present the most often on images of malignant brain tumors are P3 and P4. There are strong hopes that automatic grading can be useful during intraoperative settings and to validate effectiveness of immunotherapies [3]. Future work could provide external validation by evaluating the proposed deep learning model and quantifying protocol on a larger dataset from multiple experiments, as well as integration with clinical data by incorporating information such as patient age, gender, and other relevant medical history, with the potential to improve the model’s accuracy in predicting tumor type and prognosis.

We predict that we would be able to computationally derive features and abundances of them that are characteristic of a particular malignancy grade to classify tumors with a higher level of certainty without the necessity of confirming by genetic testing.

## 6. Acknowledgments

This work has been supported by the European Union’s Horizon 2020 research and innovation programme under grant agreement Sano no 857533, and by the International Research Agendas programme of the Foundation for Polish Science, co-financed by the European Union under the European Regional Development Fund.

We would like to thank Szymon Mazurek for his invaluable help during the code tweaking phase - with his scientific programmer’s expertise the pipeline was run efficiently on our data.

## 7. Author contributions

M.P. and A.Cr. conceptualized the study; K.W. and P.P. conducted immunohistochemical procedures and provided photos of TMAs. M.P. analyzed the data and implemented codes. M.P. and B.K. analyzed the results. M.P. wrote the original article draft. B.K., K.W., and A.Cr did the review and editing, K.W. reviewed the visualizations. The work was supervised by B.K. and A.Cr.

## 8. New or breakthrough work to be presented

In this paper, we have presented the results of a deep learning multi-class classifier as well as the discovery and learning protocol of tumor microenvironment elements used in the context of multi-grade gliomas. We obtained our results utilizing single-stained TMA samples. This approach can pave the way to automate tumor grading in clinical settings saving time for pathologists. Detecting leukocyte infiltration by immunohistochemical staining of glioma tissue microarrays may improve diagnosis and allow monitoring of the outcome of immunotherapies.

